# Non-invasive assessment of stimulation-specific changes in cerebral glucose metabolism with functional PET

**DOI:** 10.1101/2023.09.20.558617

**Authors:** Godber M Godbersen, Pia Falb, Sebastian Klug, Leo R Silberbauer, Murray B Reed, Lukas Nics, Marcus Hacker, Rupert Lanzenberger, Andreas Hahn

**Affiliations:** Department of Psychiatry and Psychotherapy, Medical University of Vienna, Austria; Comprehensive Center for Clinical Neurosciences and Mental Health (C3NMH), Medical University of Vienna, Vienna, Austria; Department of Biomedical Imaging and Image-guided Therapy, Division of Nuclear Medicine, Medical University of Vienna, Austria

**Author notes:** Correspondence to: Rupert Lanzenberger, Prof. PD MD, or Andreas Hahn, Assoc.Prof. PD PhD MSc, Department of Psychiatry and Psychotherapy Medical University of Vienna, Austria Waehringer Guertel 18-20, 1090 Vienna, Austria.

**Keywords:** brain metabolism, functional PET, fPET, quantification, percent signal change, cerebral metabolic rate of glucose (CMRGlu)

## Abstract

Functional positron emission tomography (fPET) with [^18^F]FDG allows one to quantify stimulation-induced dynamics in glucose metabolism independent of neurovascular coupling. However, the gold standard for quantification requires arterial blood sampling, which can cause discomfort for the participant and increases complexity of the experimental protocol. These constraints have limited the widespread applicability of fPET, especially in the clinical routine. Therefore, we introduce a novel approach, which enables the assessment of the dynamics in cerebral glucose metabolism without the need for an input function.

**Methods:** We tested the validity of a mathematical derivation on the basis of two independent data sets (DS). For DS1, 52 healthy volunteers (23.2 ± 3.3 years, 24 females) completed a visuo-spatial motor coordination task (the video game Tetris®) and for DS2, 18 healthy participants (24.2 ± 4.3 years, 8 females) performed an eyes-open/finger tapping task, both during a [^18^F]FDG fPET scan. Task-specific changes in metabolism were assessed with the general linear model (GLM) and cerebral metabolic rate of glucose (CMRGlu) was quantified with the Patlak plot as the reference standard. Simplified outcome parameters, such as GLM beta values of task effects and percent signal change (%SC) of both parameters were estimated. These were compared for task-relevant brain regions and on a whole-brain level.

**Results:** In general, we observed higher agreement with the reference standard for DS1 (radiotracer administration as bolus + constant infusion) compared to DS2 (constant infusion only). Across both data sets, strong correlations were found between regional task-specific beta estimates and CMRGlu (r = 0.763…0.912). Additionally, %SC of beta values exhibited excellent agreement with %SC of CMRGlu (r = 0.909…0.999). Average activation maps showed a high spatial similarity between CMRGlu and beta estimates (Dice = 0.870…0.979) as well as %SC (Dice = 0.932…0.997), respectively.

**Conclusion:** Task-specific changes in glucose metabolism can be reliably estimated using %SC of GLM beta values, eliminating the need for any blood sampling. This approach streamlines fPET imaging, albeit with the trade-off of being unable to quantify baseline metabolism. The proposed simplification enhances the applicability of fPET, allowing for widespread employment in research settings and clinical investigations.

**GRAPHICAL ABSTRACT:** 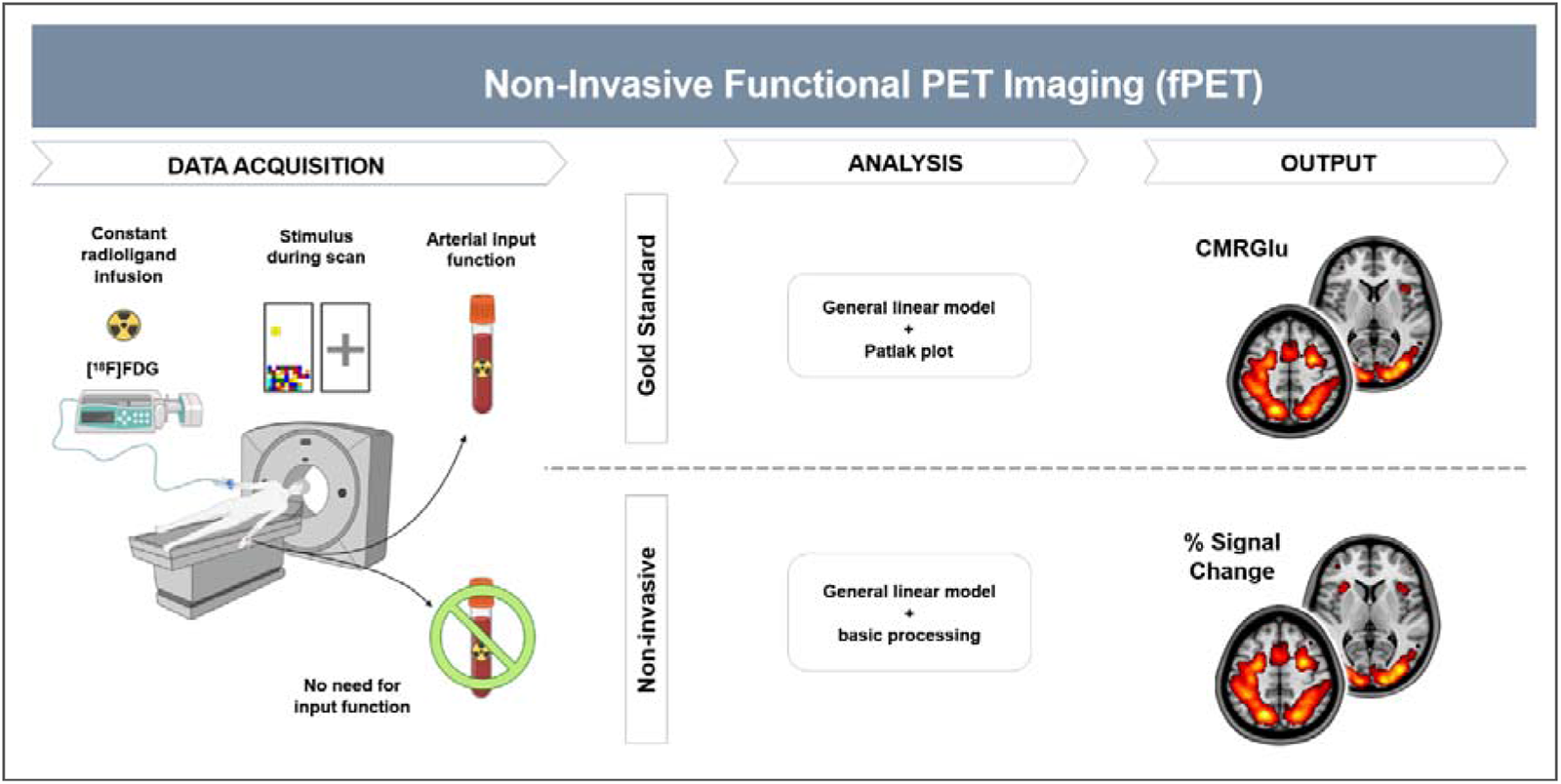

## INTRODUCTION

Functional positron emission tomography (fPET) using the radiolabeled glucose analogue 2-[^18^F]-fluorodeoxyglucose ([^18^F]FDG) holds significant promise for investigating the dynamics of brain metabolism (*1*). Using constant infusion of the radiotracer, fPET enables the assessment of changes in metabolic demands in response to external stimulation, such as cognitive tasks (*2–4*) within a single PET scan. Furthermore, these dynamics are independent from cerebral blood flow and neurovascular coupling (*2*) and the neuronal activation based on glucose metabolism can be absolutely quantified (*3*). Moreover, the widespread availability of [^18^F]FDG and the compatibility with standard PET scanners make fPET an easily accessible tool for functional neuroimaging. However, a major drawback limiting its widespread use is the need for arterial blood samples during the scan to determine the cerebral metabolic rate of glucose (CMRGlu).

The gold standard for absolute quantification in PET imaging relies on the arterial input function (AIF). However, arterial cannulation has inherent disadvantages. These include the need for skilled physicians, increased experimental complexity as well as patient discomfort or pain and in rare cases potential complications (*5*). These limitations raise the question of whether task-specific changes in glucose metabolism using fPET can be obtained without arterial blood sampling.

Several alternatives to obtain an AIF have been proposed for fPET. Venous samples (*2*,*3*,*6*) have been shown to yield sufficiently accurate quantification if the radiotracer is administered only via constant infusion (*3*). However, such a protocol results in low signal-to-noise ratio (SNR), which, among others, affects accuracy in movement correction and quantification of task effects. The use of an initial bolus resolves these issues (*7*). However, by adding a bolus, venous samples may not be adequate anymore due to a delay in the equilibration between blood pools and the subsequent underestimation of the area under the curve. The use of PBIF (*8*) is another option that avoids blood sampling. However, the assumption of equal pharmacokinetics across participants makes the approach susceptible to individual variation (*9*). Image-derived input functions (IDIF) represent another option (*10*), but robust extraction from large blood pools may be limited to total-body PET scanners. In sum, the mentioned alternatives to AIF offer easier applicability at the expense of accuracy, but may not fully eliminate the need for blood sampling.

To resolve this issue, we evaluate the feasibility of quantifying task-induced metabolic demands using [^18^F]FDG fPET without any blood sampling. We hypothesize that the input function can be omitted when task-specific activation is the primary outcome of interest. This is because the general linear model (GLM) readily separates task effects from baseline metabolism, thus yielding task-specific estimates for activation. By eliminating the requirement for blood sampling, we aim to simplify both acquisition and processing thus increasing the accessibility of fPET.

## MATERIALS AND METHODS

### Mathematical rationale

Our proposition that the GLM may be adequate for evaluating task-specific changes in glucose metabolism is grounded in the following mathematical rationale. For irreversibly binding radiotracers such as [^18^F]FDG, the ratio of tracer concentration in tissue C_T_ to that in plasma C_P_ at a certain time point t can be characterized using the Patlak plot (*11*):

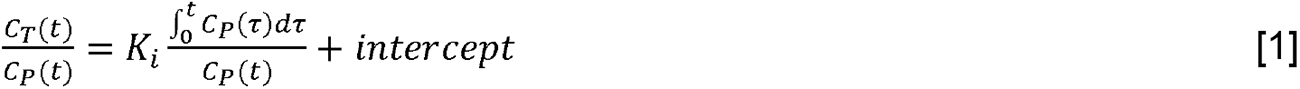

The net influx constant K_i_ is the estimated outcome parameter, which is determined as the slope of the Patlak plot when it approaches linearity after t* (*11*). The absolute amount of CMRGlu is then determined by:

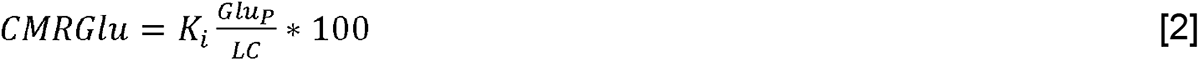

LC refers to the lumped constant and Glu_P_ represents the concentration of glucose in plasma. Given that the intercept in [1] does not change the slope of the plot (K_i_), we can therefore assume that it can be disregarded, thus rearrangement yields

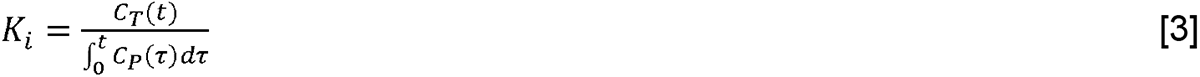

as C_P_(t) cancels out. Furthermore, in the relationship between task effects and baseline metabolism, the integral of the plasma concentration also cancels out. This implies that the ratio between the tissue concentrations is directly proportional to the relative changes in K_i_

(and thus relative changes in CMRGlu, see [2])

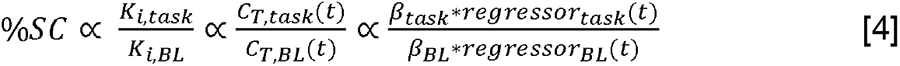

In this equation, β represents the output of the GLM when the respective regressors are used for modeling. Consequently, the ratio of the GLM’s output modeling task and baseline effects should also vary proportionally to the relative changes in CMRGlu. Since the multiplication of a beta value with its corresponding regressor represents a time course, we estimated its slope for the computation of percent signal changes (%SC, see surrogate parameters). This approach was chosen because different regressors were used for task and baseline in the GLM, which implies that simple beta values cannot be directly compared.

### In-vivo datasets

In order to test the hypothesis, we analyzed two separate datasets with different tasks and activated regions of interest (ROI). For both datasets, similar methods of preprocessing and statistical analysis were applied.

The first dataset (DS1) includes simultaneous fPET/fMRI examinations in 52 healthy participants performing a challenging visuo-spatial motor coordination task in two levels of difficulty (modified version of Tetris®). Detailed descriptions of the design, acquisition and analysis are provided in our previous work (*12*), below and in the supplement.

The second dataset (DS2) comprises data of 18 healthy participants. During the fPET/fMRI scan, participants either tapped their right thumb to their other fingers or opened their eyes. Details can be found in our previous work (*13*), below and in the supplement.

### Participants

DS1 includes 52 healthy participants (23.2 ± 3.3 years, 24 females, all right-handed), who were partly also included in previous work (*12*,*14–16*). DS2 comprises 18 healthy participants’ data (24.2 ± 4.3 years, 8 females, all right-handed), of which 15 had previously contributed to another study (*3*). See supplement for details.

### PET/MRI data acquisition and data processing

Administration of [^18^F]FDG was done according to a bolus plus constant infusion protocol for DS1 and with constant infusion only for DS2. This enables the assessment of the performance of both administration protocols. Data pre-processing of both studies’ fPET data was done with SPM12 and included motion correction, spatial normalization to MNI-space and smoothing. For both datasets manual arterial blood samples were collected to construct the AIF. See supplement for details.

### Quantification of CMRGlu

In order to analyze task activation within the two datasets, a general linear model (GLM) was applied. Both models included one regressor for baseline, one for movement artifacts and two regressors associated with task activation. For DS1, these regressors referred to the two levels of task difficulty. For DS2, they represented the separate tasks of eyes-open and right finger-tapping (see supplement).

For the calculation of the respective influx constants (K_i_), the relevant Patlak plots were constructed and their respective slopes were identified as in [1]. The start of the linear fit for the Patlak plot was set to approximately a third of the total scan time for both datasets, t*=15 min for DS1 and t*=30 min for DS2. The absolute quantification of CMRGlu was conducted in accordance with [2] and a value for the LC of 0.89, in both cases (*17*,*18*). The amount of CMRGlu was quantified in units of µmol/100g/min.

### Surrogate parameters

Our primary goal was to obtain a metric that enables the identification of task-specific changes in glucose metabolism without invasive blood sampling. Thus, we compared four different parameters of interest: i) the absolutely quantified values for CMRGlu (see [2]), used as the gold standard, ii) the plain beta values calculated by the GLM, and iii-iv) the percent signal change (%SC) of both quantities in relation to the baseline condition (see [4]). Thereby, we established a relationship between the beta values and CMRGlu as well as %SC of betas with %SC of CMRGlu. %SC for CMRGlu was calculated as the ratio of task effects to baseline metabolism multiplied by 100. The %SC for the beta values cannot be directly retrieved from the GLM output since the betas are associated with different regressors. Consequently, the slopes of the time activity curves were estimated (in kBq/frame), represented by beta*regressor separately for task and baseline metabolism (see [4]). For the baseline, a similar time interval was chosen as for the Patlak plots. For DS1 the linear fit started from minute 16 after the beginning of the radiotracer application until the end of the PET scan. For DS2, the interval began later due to the absence of an initial bolus, specifically from 30 minutes after the beginning until the end. Since the task regressors were modeled as ramp functions with a slope of 1 kBq/frame, the beta values for the tasks are already equivalent to the slope we aimed to extract. Hence, %SC of betas was then calculated as the ratio of the task and baseline slopes multiplied by 100.

Furthermore, two different baseline metrics (BL, BL2) were considered. Notably, for BL and BL2, no %SC data could be calculated, as the percent signal change inherently refers to the baseline condition itself. BL simply represents the beta value of the baseline condition as calculated by the GLM. BL2 was determined by calculating the slope of the curve given by multiplying the baseline regressor with the corresponding baseline beta values. We opted for the second baseline metric because this calculation also enters the determination of %SC of the beta values, allowing for a direct comparison. Furthermore, BL2 takes the individual variation in the baseline regressor into account and is therefore comparable across participants. It is worth noting that BL is also identical to standard uptake value ratios (SUVR) with reference to global tracer uptake. That is, regional tracer uptake is represented by regional baseline beta * baseline regressor (*3*) and since the baseline regressor represents the global tracer uptake, this cancels out when computing the ratio.

### Statistical analysis

The ROI analysis focused on the respective regions of significant activation (all p<0.05 FWE corrected) for each dataset. Linear regression analysis was performed for each pair of outcome parameters using MATLAB R2018b. For the ROI-specific analysis of DS1, the focus was placed on the frontal eye field (FEF), the intraparietal sulcus (IPS) and the secondary occipital cortex (Occ), as defined previously (*15*). DS2 displayed significant task activation in the primary occipital cortex (V1) as well as the primary motor cortex (M1) (*13*), which were used as the relevant ROI.

For the voxel-wise analysis, group-level statistics were computed in SPM12 and a one-sample t-test was performed for each of the four parameters. Activation maps were extracted (all p<0.05 FWE corrected cluster level following p<0.001 uncorrected voxel level) and activation patterns across different approaches were compared using the Dice coefficient. For DS1, we extracted respective activation maps for the hard task difficulty and for DS2 for both open eyes and right finger-tapping tasks.

## RESULTS

### Region of interest analysis

For DS1, we observed strong associations between GLM beta values and gold standard CMRGlu values for task-specific estimates of activation in the FEF, IPS and Occ (r = 0.833…0.912, Table 1). Crucially, near perfect correlations were discovered when comparing %SC of beta values with %SC of CMRGlu (all r ≥ 0.998). Moreover, the slopes were close to unity (1.00…1.02) and intercepts were near zero (-0.27…0.13) for the parameters of %SC. DS2 showed similar results for task-related changes in glucose metabolism (eyes open and finger-tapping tasks, activating V1 and M1, respectively), albeit with slightly lower performance compared to DS1. Specifically, the correlation coefficients were higher for %SC (r = 0.909… 0.970) than beta values (r = 0.763…0.833). The slopes for %SC were close to one (1.03…1.15), but intercepts were slightly higher (1.63…4.73).

**Table 1:**
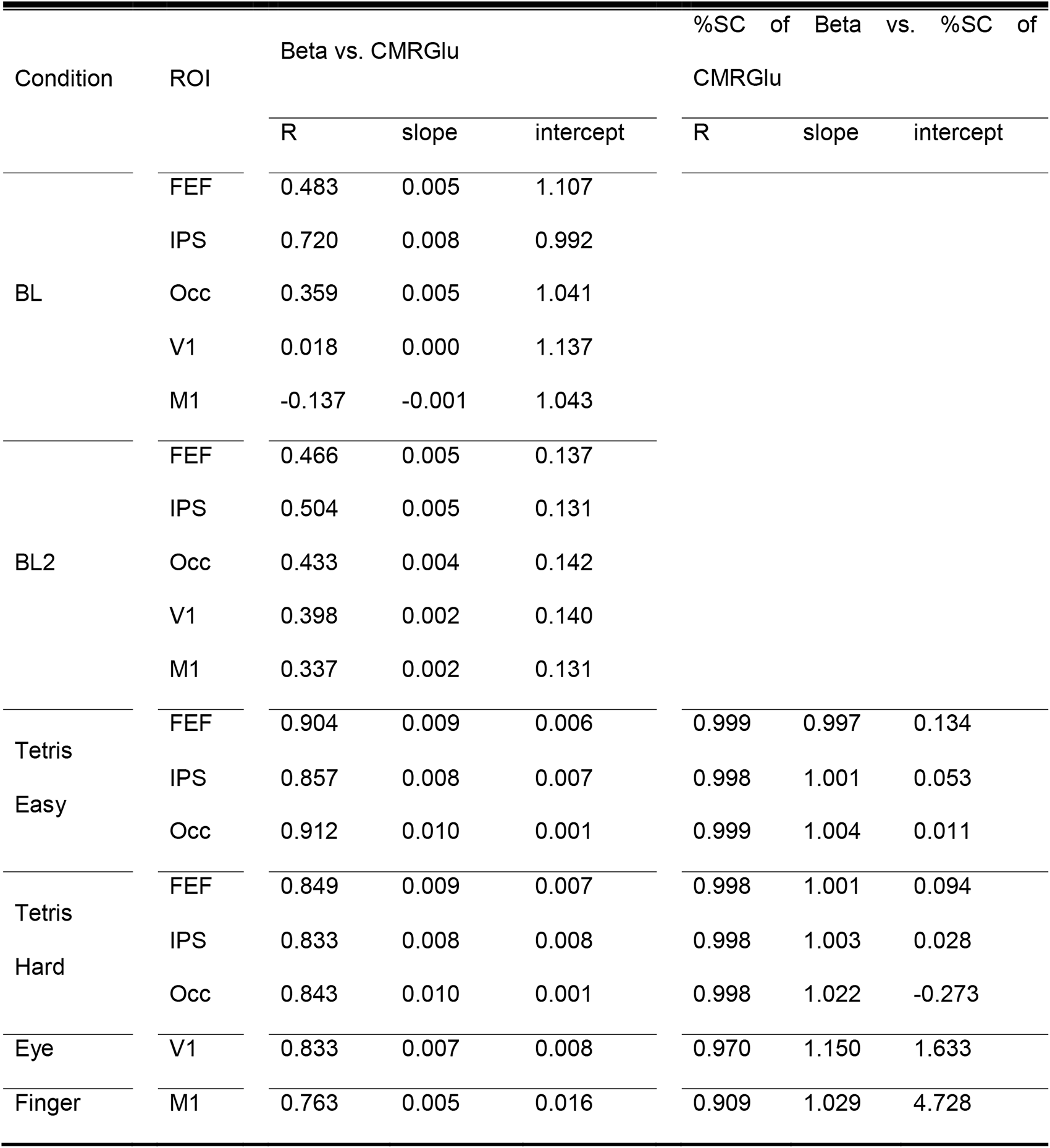
Agreement between different quantification methods. The table displays the results of correlation and regression analyses conducted for both datasets. Comparisons were performed for two different levels, either relating the GLM beta values to the respective CMRGlu (left) or the percent signal change (%SC) of both quantities with each other (right). The first dataset (DS1) comprised three regions of interest (ROI): the frontal eye field (FEF), intraparietal sulcus (IPS) and occipital cortex (Occ). For these regions, two separate levels of task difficulty (easy, hard) were regarded. For the second dataset (DS2), the primary visual (V1) and motor cortices (M1) were evaluated during the eyes-open condition and right-finger-tapping task, respectively. For all datasets, two approaches for the computation of baseline metabolism (BL and BL2) were calculated. For each comparison, Pearson’s correlation coefficient, slope and intercept were calculated. For the baseline conditions, the %SC analyses were not performed, as this parameter always refers to the baseline condition itself.

| Condition | ROI | Beta vs. CMRGlu |  |  | %SC of Beta vs. %SC of CMRGlu |  |  |
| --- | --- | --- | --- | --- | --- | --- | --- |
|  |  | R | slope | intercept | R | slope | intercept |
| BL | FEF | 0.483 | 0.005 | 1.107 |  |  |  |
|  | IPS | 0.720 | 0.008 | 0.992 |  |  |  |
|  | Occ | 0.359 | 0.005 | 1.041 |  |  |  |
|  | V1 | 0.018 | 0.000 | 1.137 |  |  |  |
|  | M1 | -0.137 | -0.001 | 1.043 |  |  |  |
| BL2 | FEF | 0.466 | 0.005 | 0.137 |  |  |  |
|  | IPS | 0.504 | 0.005 | 0.131 |  |  |  |
|  | Occ | 0.433 | 0.004 | 0.142 |  |  |  |
|  | V1 | 0.398 | 0.002 | 0.140 |  |  |  |
|  | M1 | 0.337 | 0.002 | 0.131 |  |  |  |
| Tetris<br>Easy | FEF | 0.904 | 0.009 | 0.006 | 0.999 | 0.997 | 0.134 |
|  | IPS | 0.857 | 0.008 | 0.007 | 0.998 | 1.001 | 0.053 |
|  | Occ | 0.912 | 0.010 | 0.001 | 0.999 | 1.004 | 0.011 |
| Tetris<br>Hard | FEF | 0.849 | 0.009 | 0.007 | 0.998 | 1.001 | 0.094 |
|  | IPS | 0.833 | 0.008 | 0.008 | 0.998 | 1.003 | 0.028 |
|  | Occ | 0.843 | 0.010 | 0.001 | 0.998 | 1.022 | -0.273 |
| Eye | V1 | 0.833 | 0.007 | 0.008 | 0.970 | 1.150 | 1.633 |
| Finger | M1 | 0.763 | 0.005 | 0.016 | 0.909 | 1.029 | 4.728 |

In contrast, the baseline condition (BL) displayed a highly variable degree of association with CMRGlu, with r = 0.359…0.720 for DS1 and r = -0.137…0.018 for DS2. Although BL2 resulted in more stable agreement, correlations with CMRGlu were still low (r = 0.337…0.504).

### Voxel-wise activation maps

In addition to the ROI analysis, we conducted an unbiased whole-brain analysis to explore whether the different approaches yield similar activation patterns (all p < 0.05 FWE corrected cluster level following p < 0.001 uncorrected voxel level).

As in our previous work (*12*,*15*), task-related changes in CMRGlu were observed mainly in the FEF, IPS and Occ for DS1 (Figure 2A). Interestingly, this was also true for all of the other parameters, namely maps representing beta values as well as %SC of beta and %SC of CMRGlu (Figure 2B-D). For the easy and hard levels of difficulty, the dice coefficients of the beta maps and their respective CMRGlu counterparts amounted to 0.972 and 0.979, indicating high similarity. In accordance with the ROI results, comparing the %SC maps yielded even higher dice coefficients of 0.997 (for both conditions) between the parameters.

For DS2, task-induced changes in CMRGlu occurred within V1 and M1 for the eyes open and finger-tapping tasks, respectively (*13*) (Figure 2E). Again, the activation patterns for each of the two tasks were remarkably similar across all parameters (Figure 2F-H). The dice coefficients of the beta and CMRGlu maps for the eyes open and finger-tapping tasks amounted to 0.885 and 0.870, respectively. The %SC data yielded the coefficients 0.943 and 0.932.

## DISCUSSION

In this work, we evaluated the feasibility of non-invasively quantifying task-induced changes in glucose metabolism with [^18^F]FDG fPET, i.e., without any blood sampling. We integrated theoretical concepts from the Patlak plot with output parameters of the GLM. Moreover, we compared this with CMRGlu quantified with the gold standard arterial input function in various tasks. Our findings reveal remarkably similar activation patterns across all parameters and excellent agreement between the relative changes of glucose metabolism (%SC CMRGlu) and task-specific beta values obtained from the GLM (%SC betas).

Our proposed fPET technique differs from previous approaches in one important aspect, namely its independence from an input function in general, be it arterial, venous, image-derived or population-based. The almost perfect correlation between %SC of task beta estimates and %SC of CMRGlu (i.e., r>0.998, slope∼1, intercept∼0, Table 1, Figure 1) and the virtually identical activation patterns across different parameters (Figure 2) validates the non-invasive approach as a robust alternative for performing fPET. This suggests that the underlying theoretical framework aligns excellently with the experimental data. Moreover, our approach appears to be suitable for tasks of different complexity, such as the demanding Tetris® paradigm and the simpler visual and finger tapping tasks. The slightly reduced performance in DS2 likely reflects lower SNR due to the constant infusion of the radiotracer, without the initial bolus. Nevertheless, the use of %SC is advantageous for participants as it eliminates the need for arterial cannulation and simplifies experimental procedures. This may be particularly valuable in clinical settings, where resources are often limited and procedural complexity should be minimized. Consequently, the adoption of %SC enhances the applicability of fPET in clinical environments and opens up new possibilities for diagnostic procedures beyond static PET imaging in patient cohorts (*19*).

**Figure 1:**
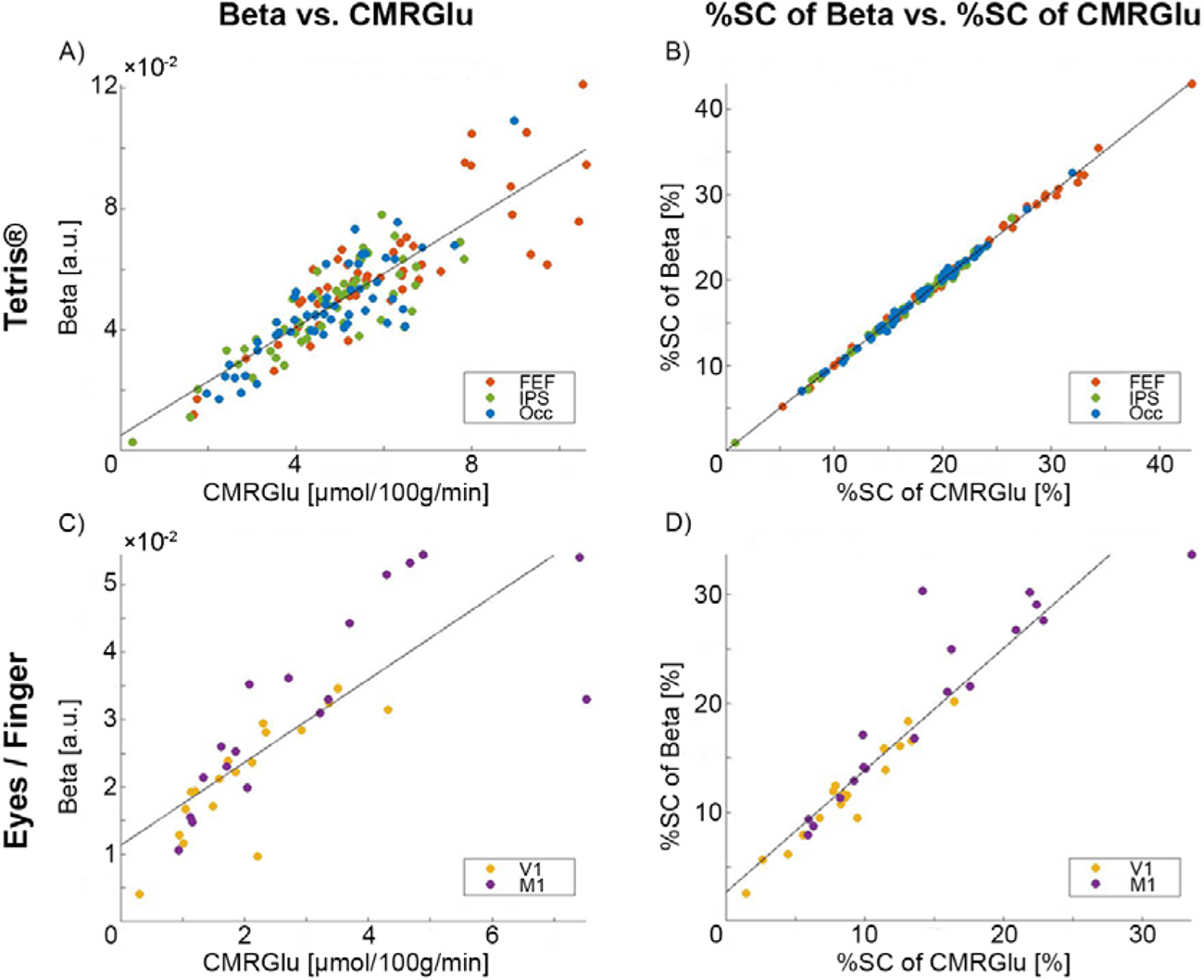
Analysis of the datasets with respect to metabolic changes in the region of interest (ROI). The figure displays the results of the regression analysis to assess whether beta values, obtained by applying the general linear model (GLM), are correlated with the cerebral metabolic rate of glucose (CMRGlu) across all participants. This was done for beta and CMRGlu values (A, C), as well as for their percent signal change (%SC) values (B, D). The figure compares these sets of analysis for task “hard”, for the Tetris®-dataset (DS1, A-B), and eye opening as well as right finger-tapping for the second PET-MR dataset (DS2, C-D). For DS1, the frontal eye field (FEF), the intraparietal sulcus (IPS) and the secondary occipital cortex (Occ) were considered as ROI. DS2 displayed activation in the primary visual cortex (V1) for eye opening and the primary motor cortex (M1) for finger-tapping.

**Figure 2:**
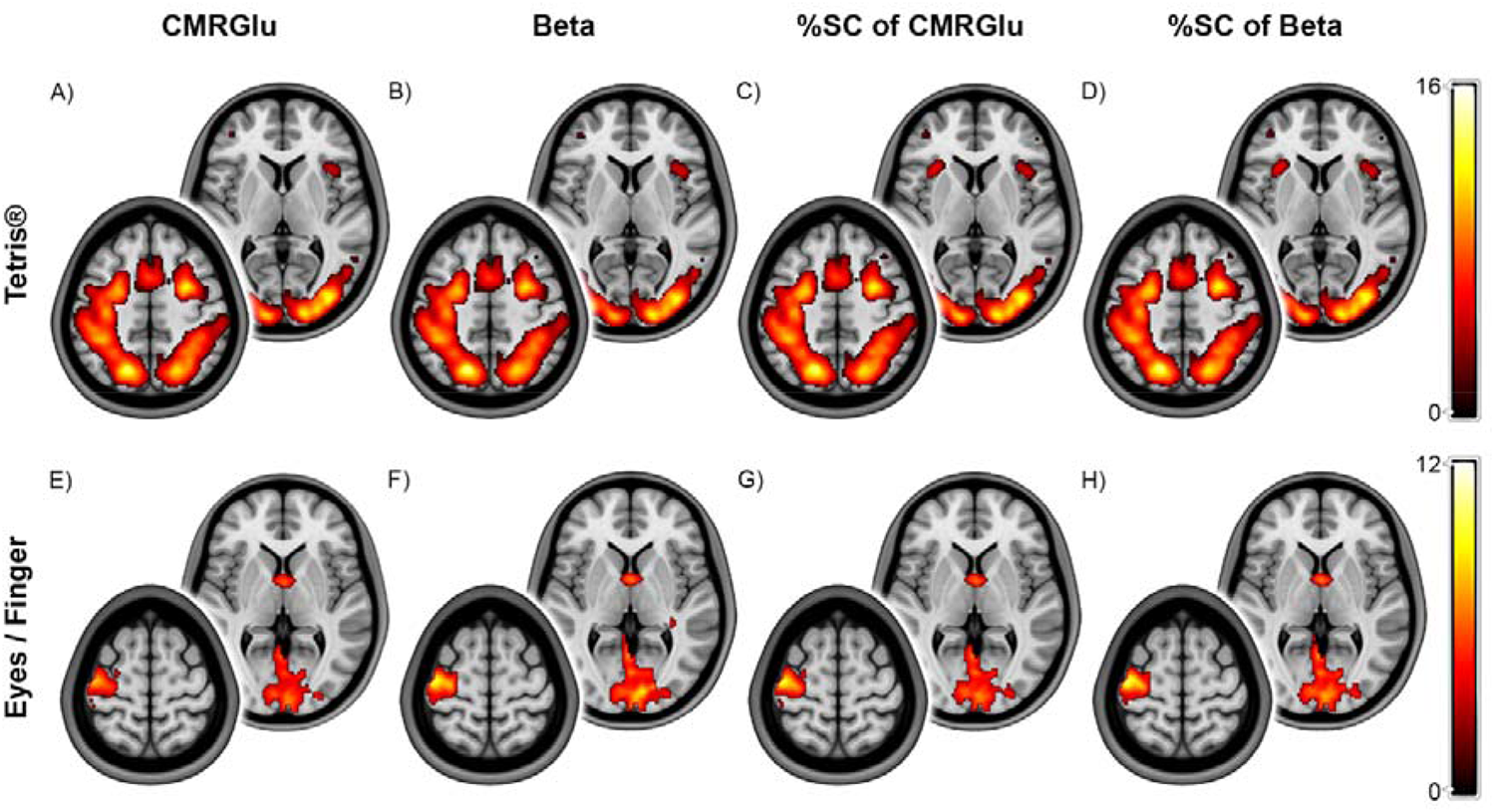
Group-level maps of the datasets, displaying activation within the respective regions of interest (ROI). The figure displays the activation patterns for both of the regarded datasets, considering task “hard” for the Tetris®-dataset (DS1, A-D), and both tasks within the second dataset (DS2, E-H). The maps were p < 0.05 FWE corrected at cluster level following p < 0.001 uncorrected voxel level. Group-level maps were calculated for the beta parameters (B, F), resulting from the general linear model (GLM), and the cerebral metabolic rate of glucose (CMRGlu, A, E) as well as for both quantities’ rate of percent signal change (%SC, C-D, G-H). For each of the group-level maps, two layers were selected to represent the activation within the respective dataset. For DS1 (A-D), the figure displays layers extracted at z = 6mm (right) and z = 50 mm (left). For DS2 (E-H), the regarded layers are z = 3mm (right) and z = 63mm (left). The colorbars represent t-values of the group level analysis.

However, it is important to acknowledge certain limitations of the simplified fPET approach (Table 2). The obtained metabolic changes are relative to a baseline condition when using %SC of betas as outcome parameter. Moreover, absolute quantification is not possible, neither for task nor baseline effects. Therefore, the technique is only suitable when the precise identification of baseline metabolism and absolute quantification are of no interest. On the other hand, the baseline definition itself becomes critical as different baselines (e.g., eyes closed, eyes open, crosshair fixation, etc.) will result in a different %SC. This reflects a similar situation encountered in fMRI, where the contrast of interest (compared to baseline or a control task) determines increases or decreases in activation (*20*,*21*). Furthermore, there is a monotonic increase of the signal over time due to the inherent property of [^18^F]FDG to remain mostly trapped in the cell. Therefore, not only the definition but also the timing of the baseline acquisition becomes relevant. To ensure robust modeling, it is recommended to acquire baseline periods in the beginning and end of the scan as well as between tasks (*7*).

**Table 2:** Visual representation of the three outcome parameters and their main features. The table displays several key features of the main outcome parameters, as obtained by [^18^F]FDG fPET and analysis with the general linear model (GLM). The parameters include the beta maps as output of the GLM, the percent signal change (%SC) of beta estimates (see [4]) and the gold standard cerebral metabolic rate of glucose (CMRGlu). The general availability of a feature for a certain outcome parameter is marked by a tick, advantages are indicated by a plus sign and disadvantages by a minus sign.

| Feature | Beta estimate | %SC of beta | CMRGlu |
| --- | --- | --- | --- |
| Identification of overall task activation | ✓ | ✓ | ✓ |
| Identification of individual effects |  | ✓ | ✓ |
| Absolute quantification |  |  | + |
| Quantification of baseline metabolism |  |  | + |
| Influence of baseline definition |  | - |  |
| Input function required |  |  | - |
| Applicable for tasks of different complexity | ✓ | ✓ | ✓ |

Regarding the use of plain beta estimates (without additional computation of %SC), it is important to note that the agreement with CMRGlu across individuals was generally lower for task effects, and poor for baseline metabolism (Figure 1, Table 1). Despite this, group-level activations still exhibited high similarity (Figure 2), suggesting that this outcome parameter may only be used to identify overall activation patterns. However, %SC only requires minimal computational effort and is thus preferable, particularly if individual values are to be related to other metrics of behavior or disease progression. Interestingly, a previous study has reported task-induced signal changes of approximately 2% (*4*), in contrast to 20-30% observed in this work (Figure 1B, 1D). However, their %SC was calculated only as a ratio of plain betas and using the same ratio for our data would result in changes in a similar range of approximately 3% (DS1). This discrepancy (and presumably also the lower agreement with CMRGlu) arises from the fact that beta values alone can be compared across participants only if the underlying regressors are identical. For this reason, we computed %SC from the slope of the product of beta*regressor (see [4]).

Although the use of fPET %SC as a proxy of neuronal activation may at first glance appear similar to BOLD fMRI, several essential differences should be kept in mind. The BOLD signal is a composite signal derived not only from neuronal oxygen consumption but also from variations in cerebral blood flow and volume (*22*), while glucose metabolism is a more direct measure of synaptic activity (*23*,*24*). Furthermore, fPET is independent of cerebral blood flow, as demonstrated by hypercapnia experiments (*2*). Thus, BOLD fMRI and [^18^F]FDG fPET capture complementary aspects of neuronal activation, as demonstrated by task-evoked dissociations between the two parameters in the default mode network (*8*,*16*,*25*). Another significant distinction lies in the test-retest variability of the methods. Previous work has indicated higher reliability for fPET than for fMRI (*14*,*26*). As a consequence, fPET seems to be a promising approach to compare intra-individual changes over time or group comparisons between imaging sites. Moreover, the approach might be relevant to assess changes in neuronal activation as induced by more potent stimulations, such as pharmacological interventions and brain stimulations.

### Conclusions

Our results suggest that plain beta estimates from the GLM may only be suitable when the overall group-averaged activation pattern is to be identified. However, computing %SC of beta values only requires minimal additional effort and represents a valid parameter to study task activation with fPET. Our data further indicates that the introduced approach is generalizable across cognitive domains and load. Still, differences between tasks may occur, which should be considered when defining the baseline condition or using control tasks for comparison. Finally, if absolute CMRGlu and baseline metabolism are of interest, full quantification is required. In sum, assessing task-specific changes in glucose metabolism with %SC is a simple and robust approach that eliminates the need for potentially painful and resource-intensive arterial blood sampling, thereby increasing the accessibility of the technique. The removal of barriers could facilitate the integration of fPET into clinical settings, where arterial blood sampling has traditionally been a major limitation.

## ACKNOWLEDGEMENTS

We thank the graduated team members and the diploma students of the Neuroimaging Lab (NIL, head: R. Lanzenberger) as well as the clinical colleagues from the Department of Psychiatry and Psychotherapy for clinical and/or administrative support. In detail, we would like to thank S Kasper, K Papageorgiou, P Michenthaler, T Vanicek, A Basaran, M Hienert, J Unterholzner and G Gryglewski, G. Karanikas for medical support, V Ritter, K Einenkel and E Sittenberger for participant recruitment and A Jelicic for partly implementation of the task. We are further grateful to J Völkle, A Pomberger, V Pichler, W Wadsak and the radioligand synthesis team from the Department of Biomedical Imaging and Image-guided Therapy, Division of Nuclear Medicine for acquisition support and supervision.

The scientific project was performed with the support of the Medical Imaging Cluster of the Medical University of Vienna.

The graphical abstract was created with BioRender.com.

## AUTHOR CONTRIBUTIONS

Study design: A.H., M.H., R.L.

Data acquisition: G.M.G., S.K., P.F., A.H., L.S, L.N.

Methods: A.H., P.F.

Data analysis: A.H., P.F., M.B.R

Manuscript preparation: G.M.G., P.F., A.H.

All authors discussed the implications of the findings and approved the final version of the manuscript

## DATA AVAILABILITY

Raw data will not be publicly available due to reasons of data protection. Processed data and custom code can be obtained from the corresponding author with a data-sharing agreement, approved by the departments of legal affairs and data clearing of the Medical University of Vienna.

