## Supplement for "Non-invasive assessment of stimulation-specific changes in cerebral glucose metabolism with functional PET"

or

Andreas Hahn, Assoc.Prof. PD PhD MSc

Department of Psychiatry and Psychotherapy

Medical University of Vienna, Austria

Waehringer Guertel 18-20, 1090 Vienna, Austria

### MATERIALS AND METHODS

#### A detailed description of the respective studies’ procedures can be found in our previous works (*1*,*2*).

#### Experimental design and tasks

The experimental design of DS1 consisted of a T1-weighted structural MRI and subsequent fPET/fMRI acquisition (52 min). After an initial baseline period (8 min), participants completed four separate sessions of Tetris® with a duration of six minutes each (2x easy and 2x hard level of difficulty, randomized). After each task block, an additional resting period of five minutes was scheduled. The task for DS1 was a modified version of Tetris®. The levels of difficulty were defined by the speed of the falling bricks and the already existing number of bricks at the start of each round. During the task, participants were able to control the game with their right hand only. All participants underwent an initial training, in which they practiced the task in both difficulties for 30 seconds each, before the scan. All periods of rest were defined by participants looking at a crosshair and letting their thoughts wander freely. Regarding DS1, only the data of the first PET/MR measurement (*3*), was analyzed in this work.

For the collection of the DS2, participants were examined in a 95-minute-long fPET/fMRI scan, which included four separate ten-minute blocks of task performance. Between each of these task blocks, a 15-minute period of rest was scheduled. At the beginning of the scan, a T1-weighted structural image was recorded. The four blocks of task performance consisted of participants opening their eyes at 10-20 and 60-70 min as well as tapping their right thumb to their fingers at 35-45 and 85-95 min after the beginning of the radiotracer administration. During the initial baseline and the periods of rest, participants had their eyes closed and did not move their fingers.

#### Participants

For both datasets, participants underwent medical pre-examinations as well as the Structural Clinical Interview for DSM-IV, led by an experienced psychiatrist. These also included an assessment of the general state of health, blood laboratory tests, electrocardiography and a neurological evaluation. Exclusion criteria were former or current somatic, neurological or psychiatric disorders, former or current substance abuse and medication intake, previous study-related radiation exposure as well as pregnancy or breastfeeding. For DS1, specifically, it was also essential that participants did not have any previous experience playing Tetris® within the past three years.

All participants gave their written informed consent for participation after extensive instruction about the respective study’s protocol. Both studies were approved by the Ethics Committee of the Medical University of Vienna (ethics numbers DS1: 1479/2015 and DS2: 1916/2013). Procedures were carried out in agreement with the Declaration of Helsinki. The study related to DS1 has been registered at ClinicalTrials.com (clinical trials identifier: NCT03485066).

#### PET/MRI data acquisition and data processing

DS1 participants were administered with [^18^F]FDG according to a bolus plus constant infusion protocol (510 kBq/kg/frame for 1 min and 40 kBq/kg/frame for 51 min). This was realized by employing a perfusion pump (Syramed µSP6000, Arcomed, Regensdorf, Switzerland). The PET data comprised by DS1 was corrected for attenuation and reconstructed using the ordinary Poisson ordered subset expectation maximization algorithm (OP-OSEM) (3 iterations, 21 subsets) into 30-second frames, with an overall matrix size of 344 x 344 and 127 slices. The structural MRI was recorded by applying a T1-weighted MPRAGE sequence (TE/TR = 4.21/2200 ms, voxel size = 1 x 1 x 1.1 mm, 7.72 min).

For DS2, the radiotracer was administered according to a constant infusion protocol which lasted the entire scan. The dose was 3 MBq/kg bodyweight and the radiotracer was distributed at a speed of 36 ml/h by a pump (Volumed µVP7000, Arcomed, Regensdorf, Switzerland). The PET data was reconstructed into one-minute frames using OP-OSEM and corrected for attenuation via an additional CT or a pseudo-CT (*4*), calculated from a T1-weighted image, for one participant. The T1-weighted structural image was obtained by a MPRAGE sequence (TE/TR = 4.2/2000 ms, voxel size = 1 x 1 x 1.1 mm).

Data pre-processing of both studies’ fPET data was done using SPM12 and included motion correction (quality = 1, registered to mean), spatial normalization to MNI-space with transformation matrices obtained from the structural MRI and smoothing with an 8 mm Gaussian kernel. Masking of the datasets led to the exclusion of non-gray-matter voxels and low-pass filters were used. The cutoff frequency was set to half of the task duration, i.e., 3 min for DS1 and 5 min for DS2.

#### Blood sampling

For both datasets manual arterial blood samples were collected. For DS1, this was done at 3, 4, 5 min as well as 14, 25, 36, 47 min after the start of the tracer application. For DS2 arterial samples were taken at 10, 20, 35, 45, 60, 70, 85 and 95 min after the start of the radiotracer application. The respective arterial samples of both datasets were analyzed regarding their whole-blood and plasma activity with a γ-counter (Wizward2, 3”, Perkin Elmer), which was cross-calibrated to the PET/MR scanner. This enabled the construction of individual AIFs while correcting for the plasma-to-whole-blood ratio. For DS1, this was realized by linear interpolation (*5*), while data of DS2 were modeled with the sum of two exponential functions.

#### Quantification of CMRGlu

The task regressors were modeled as linear ramp functions with a slope of 1 kBq/frame. In the case of DS1, the baseline regressor was calculated by averaging the time course across all gray matter voxels, but excluding those voxels that were declared as active by the fMRI (p < 0.05 FWE corrected voxel level) (*5*). For DS2, the baseline regressor was estimated by averaging the time course across all gray matter voxels, which was modeled by a third order polynomial function while controlling for task effects. Although these approaches slightly differ, we have previously shown that the results are highly comparable and do not affect test-retest reliability (*1*,*5*,*6*). Finally, the regressor accounting for movement artifacts was defined by the first principal component of the six realignment parameters, for both datasets.
